# Re-curation and Rational Enrichment of Knowledge Graphs in Biological Expression Language

**DOI:** 10.1101/536409

**Authors:** Charles Tapley Hoyt, Daniel Domingo-Fernández, Rana Aldisi, Lingling Xu, Kristian Kolpeja, Sandra Spalek, Esther Wollert, John Bachman, Benjamin M. Gyori, Patrick Greene, Martin Hofmann-Apitius

**Affiliations:** Department of Bioinformatics, Fraunhofer Institute for Algorithms and Scientific Computing (SCAI), Sankt Augustin 53754, Germany; Bonn-Aachen International Center for IT, Rheinische Friedrich-Wilhelms-Universität Bonn, Bonn 53115, Germany; Laboratory of Systems Pharmacology, Harvard Medical School, 200 Longwood Ave, 02115 Boston, MA, USA

**Keywords:** Natural language processing, Information extraction, Biocuration, Biological Expression Language, Knowledge graphs

## Abstract

The rapid accumulation of new biomedical literature not only causes curated knowledge graphs to become outdated and incomplete, but also makes manual curation an impractical and unsustainable solution. Automated or semi-automated workflows are necessary to assist in prioritizing and curating the literature to update and enrich knowledge graphs.

We have developed two workflows: one for re-curating a given knowledge graph to assure its syntactic and semantic quality and another for rationally enriching it by manually revising automatically extracted relations for nodes with low information density. We applied these workflows to the knowledge graphs encoded in Biological Expression Language from the NeuroMMSig database using content that was pre-extracted from MEDLINE abstracts and PubMed Central full text articles using text mining output integrated by INDRA. We have made this workflow freely available at https://github.com/bel-enrichment/bel-enrichment.

**Database URL:** https://github.com/bel-enrichment/results

## Background

The rapid accumulation of unstructured knowledge in the biomedical literature has motivated its structuring and formalization so computers can assist in large-scale reasoning and interpretation. Several standard formats have been proposed for storing newly structured knowledge, including Systems Biology Markup Language (SBML; Hucka *et al*., 2003), Biological Pathways Exchange Language (BioPAX; Demir *et al*., 2010), Biological Expression Language (BEL; Slater, 2014), Gene Ontology Causal Assembly Models (CAMs; Carbon *et al*., 2017). Accompanying these standards are public repositories containing content generated both in academic and industrial contexts such as the BioModels Database (Glont *et al*., 2018), Pathway Commons (Cerami *et al*., 2011), NDEx (Pratt *et al*., 2015), Bio2RDF (Belleau *et al*, 2008), Open PHACTS (Williams *et al*, 2012), and BEL Commons (Hoyt *et al*, 2018). Additionally, a significant number of databases use custom formats for knowledge that are not appropriate for formalization in a standard format.

Even though each standard focuses on different aspects of modeling knowledge in systems and networks biology, they all give rise to knowledge graphs (KGs) consisting of biological entities (nodes), their interrelations (edges), and their associated metadata. While KGs have been useful for qualitative modeling of biochemical networks (Rausanu *et al*., 2015; Yugi *et al*., 2016), cellular signaling (Pilalis *et al*., 2015; Pon *et al*., 2015; Tripathi *et al*., 2015), gene regulatory pathways and genetic interactions (Kandasamy *et al*., 2010; Kamburov *et al*., 2013), metabolic pathways (Caspi *et al*., 2016; Wishart *et al*., 2018), and other systems biology applications, there are several challenges associated with their use. First, they contain noise arising from curation, from the loss of information due to representation, and from normalization of different knowledge representations (Nickel *et al*., 2016; Mihindukulasooriya *et al*., 2017; Pujara *et al*., 2017). Second, they are generally an incomplete representation of the current state of scientific knowledge due to the large amount of uncurated, unstructured knowledge in the literature. Third, they progressively become out-of-date as scientific experimentation and investigation elucidates new knowledge (Wadi *et al*., 2016). Finally, they often lack biological contextual information such as organelle-, cell-, cell line-, tissue-, organ-, phenotype-, or disease-specificity (Hofmann-Apitius *et al*., 2015; Saqi *et al*., 2018).

KGs also suffer from issues in the normalization and mapping of entities. Though interoperability standards and resources like the Minimal Information Required in the Annotation of Models (MIRIAM; Laibe *et al*., 2007) and Identifiers.org (Juty *et al*., 2012) have been developed and implemented to promote the semantic interoperability of biological models (and by extension, KGs), curators often encounter concepts that are not present in high-quality, publicly available terminologies and can not capture the incident knowledge in a semantically meaningful way. These situations require enriching previously existing terminologies or, in some cases, developing new ones. For situations when the appropriate concept/term is unclear, several tools have been developed and made freely available to the community to help curators build semantically interoperable models including the Ontology Lookup Service (OLS; Cote *et al*., 2007), the Ontology Mapping Service (OxO; https://www.ebi.ac.uk/spot/oxo), Zooma (https://www.ebi.ac.uk/spot/zooma), and CEDAR Workbench (Gonçalves *et al*., 2017). Further, recent work from Domingo-Fernández *et al*. on mapping pathways between major databases (Domingo-Fernández *et al*., 2018) and a critical assessment of their overlaps and contradictions (Domingo-Fernández *et al*., 2019) has shown that that the adoption of standards like MIRIAM has been slow and that while the syntax of the varying formats used by each database may be correct, their semantic interoperability is still lacking.

## Motivation

Accurately structuring and formalizing the unstructured knowledge in the biomedical literature requires careful planning and manual effort from trained curators. The scope of a given project must be defined based on its scientific goals (e.g., to support the interpretation of data, to generate a disease-specific knowledgebase, etc.) and limited in its literature content sources (e.g., abstracts, full text, patents, etc.) based on a project-specific metric for quality and relevance — both of which are nebulous in description and difficult to generate. The scope must also be limited to certain classes of biological entities, their interrelations, and the standard formats that are capable of expressing them. For instance, the entities, relations, and formats used during curation are different for protein complex assemblies curated by the Complex Portal (Meldal *et al*., 2015) and regulatory interactions curated by the Signaling Network Open Resource (SIGNOR; Perfetto *et al*., 2016). Similarly, curation guidelines must be defined reflecting these limits. For example, the guidelines of a project designed to model Tau aggregation inhibitors from the chemistry literature might encourage the curators to include direct binding partners of those inhibitors (e.g., GSK-3β, CDK5, etc.) but explicitly exclude the biological mechanisms through which the inhibitors’ targets result in Tau aggregation that would better be curated during a different project focusing on capturing molecular biology from its primary literature. While there is no alternative to proper planning, several semi-automated curation workflows such as BELIEF (Madan *et al*., 2016) and the sbv IMPROVER (Guryanova *et al*., 2017) provide assistance by automatically detecting entities and relations for curators to accept or fix in order to increase productivity and enforce correct syntax and semantics. However, these and similar systems are limited in their ability to capture the relevant chemistry and biology, and reversion to manual curation is often necessary. Finally, the issues of insufficient resources and fixed timelines apply to most curation projects, as aptly described by Rodríguez-Esteban (2015).

In the AETIONOMY project (https://www.aetionomy.eu), we manually curated NeuroMMSig, an inventory of multiscale and multimodal knowledge graphs that capture mechanistic knowledge in the context of neurological disorders (Domingo-Fernández *et al*., 2017). We encoded it in BEL because it is appropriate for qualitative causal, correlative, and associative relationships between biological entities, processes, and measurements across modes and scales. However, it is currently suffering from the issues we have previously described: it has not been assessed for confidence, is becoming outdated, and needs to be enriched following a rational approach that best prioritizes the flood of recent literature.

To address this, we have developed and applied two workflows, described in this paper: the first is for re-curating existing BEL documents to ensure their syntactic and semantic correctness in a scenario where there was neither prior syntax validation, curation guidelines for entity nomenclature, nor a second curator for achieving inter-annotator agreement. The second is a semi-automated algorithm and reproducible workflow for updating and rationally enriching an existing KG that lessens the burden of identifying relevant literature, reduces the overhead, as defined by Rodríguez-Esteban, and generates more, higher quality, relevant content.

We applied these workflows to a selection of knowledge graphs in NeuroMMSig and evaluated the curation effort (time) and quality in comparison to purely manual curation and other previously reported semi-automated curation workflows. We increased the number of nodes and edges in the selected knowledge graphs respectively by approximately five and seven times while maintaining the specificity of the knowledge graphs. With an improvement to the content underlying NeuroMMSig, the mechanism enrichment algorithm on its corresponding web service can return more correct and robust results to support the analysis of neuroimaging and genomics data for clinical trials in Alzheimer’s disease, Parkinson’s disease, and epilepsy. Finally, we have made this workflow freely available at https://github.com/bel-enrichment/bel-enrichment so others can include it in their own curation workflows.

## Methods

We first present the re-curation workflow for syntactic and semantic quality assurance before presenting our proposed approach for updating and rational enrichment.

### Syntactic Quality Assurance

We developed a workflow using git (https://git-scm.com), GitHub (https://github.com), PyBEL (Hoyt *et al*., 2017), and a novel PyBEL extension PyBEL-Git (Hoyt, 2018) in order to identify and address syntactical issues in the BEL documents generated during the AETIONOMY project (https://www.aetionomy.eu: Irin *et al*., 2015; Kodamullil *et al*., 2015; Naz *et al*., 2016; Emon *et al*., 2017; Hoyt and Domingo-Fernández *et al*., 2018) and exposed through the NeuroMMSig mechanism enrichment server (Domingo-Fernández *et al*., 2017).

This workflow can be implemented in other web-based version control systems such as GitLab (https://gitlab.com) and Atlassian BitBucket (https://bitbucket.org) as well as directly integrated with continuous integration systems such as GitLab CI/CD (https://docs.gitlab.com/ee/ci). Travis-CI (https://travis-ci.com), and BitBucket Pipelines (https://bitbucket.org/product/features/pipelines) using the instructions provided at https://github.com/pybel/pybel-git with minimal configuration.

### Semantic Quality Assurance

We selected ten signatures (and their corresponding BEL documents) from NeuroMMSig based on their druggability (number of proteins targeted by drugs that have been assessed in clinical trials), their novelty (less preference given to subgraphs corresponding to hypotheses that have repeatedly failed in the clinic; namely amyloid-beta aggregation), and their amenability to assay development (based on expert advice) as an example for the re-curation workflow outlined below. An enumeration and statistics can be found in **Table 1** and the signatures can be explored through BEL Commons (Hoyt *et al*., 2018).

**Table 1:** Statistics for the number of BEL nodes and BEL statements in the ten knowledge graphs selected from the NeuroMMSig inventory before re-curation (using the version last updated on December 6^th^, 2016), after-recuration, and after enrichment. Later, we discuss these statistics in terms of INDRA statements - the discrepancies are due to the ontological reasoner applied in the conversion process from INDRA statements to BEL statements.

| Label | Description | Before Re-curation |  | After Re-curation |  | After Enrichment |  |
| --- | --- | --- | --- | --- | --- | --- | --- |
|  |  | Nodes | Edges | Nodes | Edges | Nodes | Edges |
| Tau protein subgraph | The downstream effects of the post-translational modification, aggregation, and transport of the Tau protein | 191 | 493 | 261 | 733 | 708 | 2054 |
| DKK1 Subgraph<br>GSK3 Subgraph | The interaction partners with GSK-3β and its targets of post-translational modification. The complementary DKK1 pathway is a specific signaling cascade upstream of GSK-3β | 128 | 254 | 174 | 377 | 376 | 1165 |
| Inflammatory Response | Processes related to inflammation in the context of Alzheimer's disease | 182 | 373 | 341 | 743 | 2003 | 7607 |
| Insulin Signal Transduction | The molecular relationships between insulin resistance and inflammation, motivated by epidemiological studies that suggested a correlation between AD and Type II diabetes (Karki *et al.*, 2017). | 251 | 739 | 315 | 881 | 612 | 1973 |
| Amyloidogenic Subgraph | The downstream effects of the amyloid precursor protein (APP), its protein modifiers, and its cleavage products | 493 | 1223 | 652 | 1751 | 2090 | 7436 |
| Non-amyloidogenic Subgraph | Chemicals and processes known to down-regulate the expression of the transcript corresponding to APP or the abundance of the APP protein | 195 | 359 | 325 | 635 | 795 | 2238 |
| Apoptosis and Cell Death | Processes relevant to AD that result in apoptosis including the Caspase subgraph, XIAP subgraph, and Complement system subgraph | 104 | 143 | 170 | 229 | 1065 | 2401 |
| Acetylcholine Subgraph | Pathways including biological entities and processes related to cholinergic neurons and acetylcholine transmission | 106 | 197 | 148 | 337 | 423 | 1275 |
| GABA Subgraph | Pathways including biological entities and process related to GABAergic neurons and GABA transmission | 21 | 30 | 91 | 190 | 305 | 721 |
| Reactive Oxygen Species Subgraph | The effects of reactive oxygen species, including the Myeloperoxidase subgraph, Hydrogen peroxide subgraph, Free radical formation subgraph, and Nitric oxide subgraph | 104 | 173 | 126 | 224 | 1401 | 6277 |
| **Total** |  | **1188** | **3529** | **1704** | **5391** | **5850** | **23811** |

Because BEL was developed by the biomarker discovery company, Selventa, before the wide adoption of semantic resources like Identifiers.org, the Open Biomedical Ontology (OBO) Foundry, and the OLS, the language used a custom format for storing the names and identifiers of entities in major biomedical databases and ontologies such as the HUGO Genome Nomenclature Consortium (HGNC; Yates *et al*., 2017) Chemical Entities of Biological Interest (ChEBI; Hastings *et al*., 2013), the Gene Ontology (GO; Carbon *et al*., 2017), Medical Subject Headings (MeSH; Rogers, 1963), the Disease Ontology (DO; Schriml *et al*., 2018), the Human Phenotype Ontology (HPO; Köhler *et al*., 2018), the Cell Line Ontology (CLO; Sarntivijai *et al*., 2014), the Experimental Factor Ontology (EFO; Malone *et al*., 2010), and others. Additionally, Selventa provided several entity type-specific, manually curated terminologies for chemicals, protein families, protein complexes, and diseases for entities that had not yet been included in any of the other existing resources.

Because the Selventa terminologies are no longer maintained and the publicly available terminologies have far surpassed them in coverage, the first step in re-curation was to normalize entities to high-quality, publicly available terminologies. For example, chemicals were normalized to identifiers from ChEBI, ChEMBL (Gaulton *et al*., 2017), and PubChem (Kim *et al*., 2016) whenever possible; protein families and complexes were normalized to FamPlex (Bachman *et al*., 2018); and diseases were normalized to DO and HPO. Further, because the BEL documents from AETIONOMY were all produced before 2015, the entities that were curated using their labels (instead of stable identifiers) needed to be updated. A short investigation showed that HGNC and GO were the least stable namespaces, but combined they had less than one hundred entities to be addressed. We therefore concluded that manual intervention was more appropriate than developing complicated systems for updating labels. While it is not intended to be the focus of this article, we have also begun to build a custom terminology (available at https://github.com/pharmacome/terminology) to supplement the publicly available ones for a small number (less than 1000) of terms that had not been included in other resources.

After ensuring both the correctness of BEL syntax and namespace usage, a remaining major aspect of re-curation is to address the issues arising from curation lacking inter-annotator agreement. BEL statements and their corresponding annotations (metadata) were generated by several independent curators and had not undergone quality control either by comparison with the results of independent curation of the same document by a second curator, or even minimally checked by a second curator. We applied the following simple guidelines:

1. *Second Curator*: check and label all relevant statements with a `SET Confidence` annotation using the Likert scale as described in **Table 2**.
2. *Third Curator (curation leader)*: after all relevant statements had been checked for correctness, check all statements with `SET Confidence` = “`High`” or `SET Confidence` = “`Medium`”. Change the confidence to `SET Confidence` = “`Very High`” on agreement. Otherwise, fix the statement.

**Table 2.** Confidence annotations using the Likert scale for re-curation

| Confidence | Rationale |
| --- | --- |
| None | If the evidence string is nonsense or contains no reasonable biological knowledge, delete it and the related statements entirely. It's okay to remove BEL statements that are not supported. |
| Low | If it's not clear what BEL should represent the biology, add SET Confidence = "Low" for later discussion. |
| Medium | If the statement is wrong, fix it and add the annotation SET Confidence = "Medium". |
| High | If statement can be asserted from the given evidence, add the annotation SET Confidence = "High". |

The existence of the confidence guideline can be checked with the PyBEL command line interface with the following command: `pybel compile --required-annotations` “`Confidence`”.

### Proposed Approach for Updating and Rational Enrichment

Next, we developed and applied a procedure for enriching a given BEL document in order to cope with the mounting issues of out-of-dateness and incompleteness. Our approach identifies nodes with low information density and uses a large-scale corpus of biomedical literature that has been pre-processed by automated relation extraction methods to identify the most relevant literature, evidences, and ultimately relations. Notably, the previously described quality assurance (i.e., re-curation) workflows for checking and addressing the syntactic and semantic correctness of a given BEL document were necessary to decrease the noise input into the procedure. Following the re-curation of the ten NeuroMMSig subgraphs, we applied the following procedure for rational enrichment:

1. *Knowledge Graph Pre-processing*: nodes corresponding to the same gene (i.e., RNA, microRNA, Protein, and variants thereof) are collapsed, non-causal relationships (e.g., correlative, associative, ontological, etc.) are removed, and several entity types (i.e., abunances, reactions, pathologies, and biological processes) are removed.
2. *Application of Information Density Metric*: the remaining nodes are ranked by an information density function. We used the sum of the node in-degree and out-degree as this corresponds to the amount of causal information for a given gene in the knowledge graph. In this scenario, isolated nodes correspond to genes for which there is no causal information about its interactions with other proteins, and leaves (i.e., entities with only one edge) correspond to nodes that have very limited information.
3. *Automated Relation Extraction*: the top-ranked genes are used as a query to a knowledge graph generated by large-scale automated biological relation extraction. We used the Integrated Network and Dynamical Reasoning and Assembler (INDRA; Gyori *et al*., 2018) and applied several filters to find the most relevant and novel relations. First, the relations that were already curated and in the knowledge graph were excluded. Second, INDRA was used to calculate a confidence score (between 0.0 and 1.0) for each relation based on evidences from structured databases and the frequency of occurrence of similar statements. Those statements with a low confidence score (< 0.80) were removed to increase the precision and therefore reduce the curation overhead. While INDRA integrates relations extracted from multiple reading systems, a corpus of relations from a single machine reading system, such as EVEX, would serve the same purpose (Van Landeghem *et al*., 2012).
4. *Conversion to BEL*: different automated relation extraction systems present various information (e.g., entity offsets, events, triggers, etc.) in ways that are not amenable to curation. Because INDRA already normalizes this information for several systems to several varieties of the `indra.Statement` Python class, we developed a converter to BEL using PyBEL that can be used directly with the `indra.assemblers.PybelAssembler` Python class. Finally, this information is exported to an Excel sheet with several additional columns for tracking INDRA statement provenance, curator provenance, the correctness of BEL statements, the type of errors found, and the changes made to incorrect BEL statements. Examples and links to the full results can be found in the supplementary information.

For each round of rational enrichment, the procedure was applied to generate several curation sheets corresponding to the lowest information genes. Each row was checked with the following procedure:

1. Place an “x” in the Checked column.
2. If the BEL statement correctly corresponds to the *Evidence* column, place an “x” in the Correct column.
3. Else if the BEL statement can be improved (e.g., assignment of entity types, relation, etc.), correct it and place an “x” in the *Changed* column and annotate the error type in the *Error Type* column using a controlled vocabulary (see the supplementary data). Additional guidelines for categorizing error types can be found at https://github.com/pharmacome/curation/blob/master/indra-errors.rst.
4. Else if the BEL statement does not correspond to the *Evidence* column and can not be improved, then “x” should neither be placed in the *Correct* nor the *Changed column*.
5. If the *Evidence* column contains other BEL statements that were not extracted, duplicate the current row’s provenance (reference, evidence, etc.) and add the additional BEL statements. Place an “x” in the *Changed* column but not the *Correct* column.
6. If there are other BEL statements that can be extracted, make a new line with all of the same provenance information (uuid, reference, evidence, etc.) and just place an “x” in the “Changed” column.

This procedure was applied iteratively: as the low information density nodes from the first round gained new relations, the knowledge graph was expanded and further low information density nodes were added. There are several improvements that could be made to the information density function and prioritization of the resulting extracted statements. For example, relations found by INDRA between low information density nodes and high information density nodes could be prioritized to maintain the scope and focus of a knowledge graph.

## Results and Discussion

While applying the re-curation workflow outlined in **Figure 1**, we identified large sections of poor quality curation that had to be removed. Additionally, some evidences in the BEL document that were previously incompletely curated were completed. Re-curation also required the updating of namespaces from the 2015 versions to the most current and necessitated some additional revisions.

**Figure 1:**
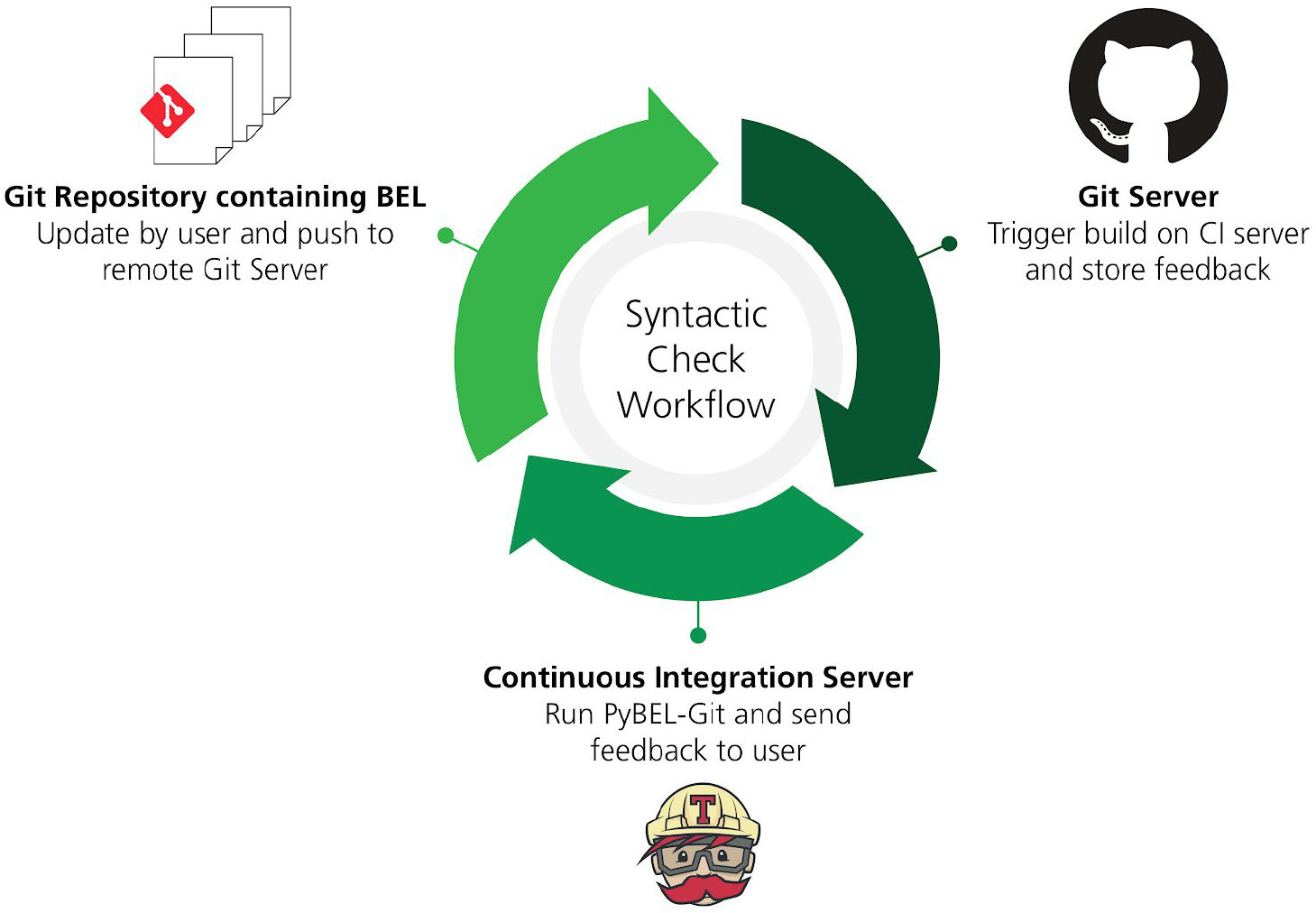
A workflow for syntactic quality assessment. This figure can be found on FigShare at https://doi.org/10.6084/m9.figshare.7643006.v1.

To evaluate the enrichment workflow outlined in **Figure 2**, we defined weekly curation rounds in which each of the five curators were tasked to curate the enrichment template generated by INDRA for the first 30 prioritized genes. Curators worked 10 hours per round for one month (4 weeks; one round per week) to curate BEL statements from a pool of 113 genes. A database of statements was generated by INDRA using the REACH (Valenzuela-Escárcega *et al*., 2015; Valenzuela-Escárcega *et al*., 2018), and Sparser (McDonald, 2000) readers to extract a total of 17096 statements containing these genes from all MEDLINE abstracts and PubMed Central full text articles available in August 2018. Of these, 2989 were manually evaluated. 917 statements (30.7%) were marked as correct by the curators, 1454 statements (48.6%) required manual corrections, and the remainder (20.7%) could not be corrected. The criteria for correctness was that *all* aspects of the statement, including the subject and object entities, relationship type, phosphorylation and other post-translational modifications, were extracted to the same extent as careful manual curation could. Ultimately, excluding the statements that could not be corrected, 79.3% of the automatically extracted, manually revised BEL statements were recovered. After curation, the recovered statements were converted into a BEL knowledge graph that contained 4228 nodes and 17002 edges complementary to the original ten subgraphs selected from NeuroMMSig. The discrepancies in the number of INDRA statements to BEL statements is due to the ontological reasoning process that occurs during conversion. For example, INDRA statements about protein complex formation are converted to bi-directional BEL statements, INDRA statements about post-translationally modified proteins induce edges to the reference protein, and INDRA statements about bound proteins create a variety of additional BEL nodes representing their constituents and membership edges connecting them.

**Figure 2.**
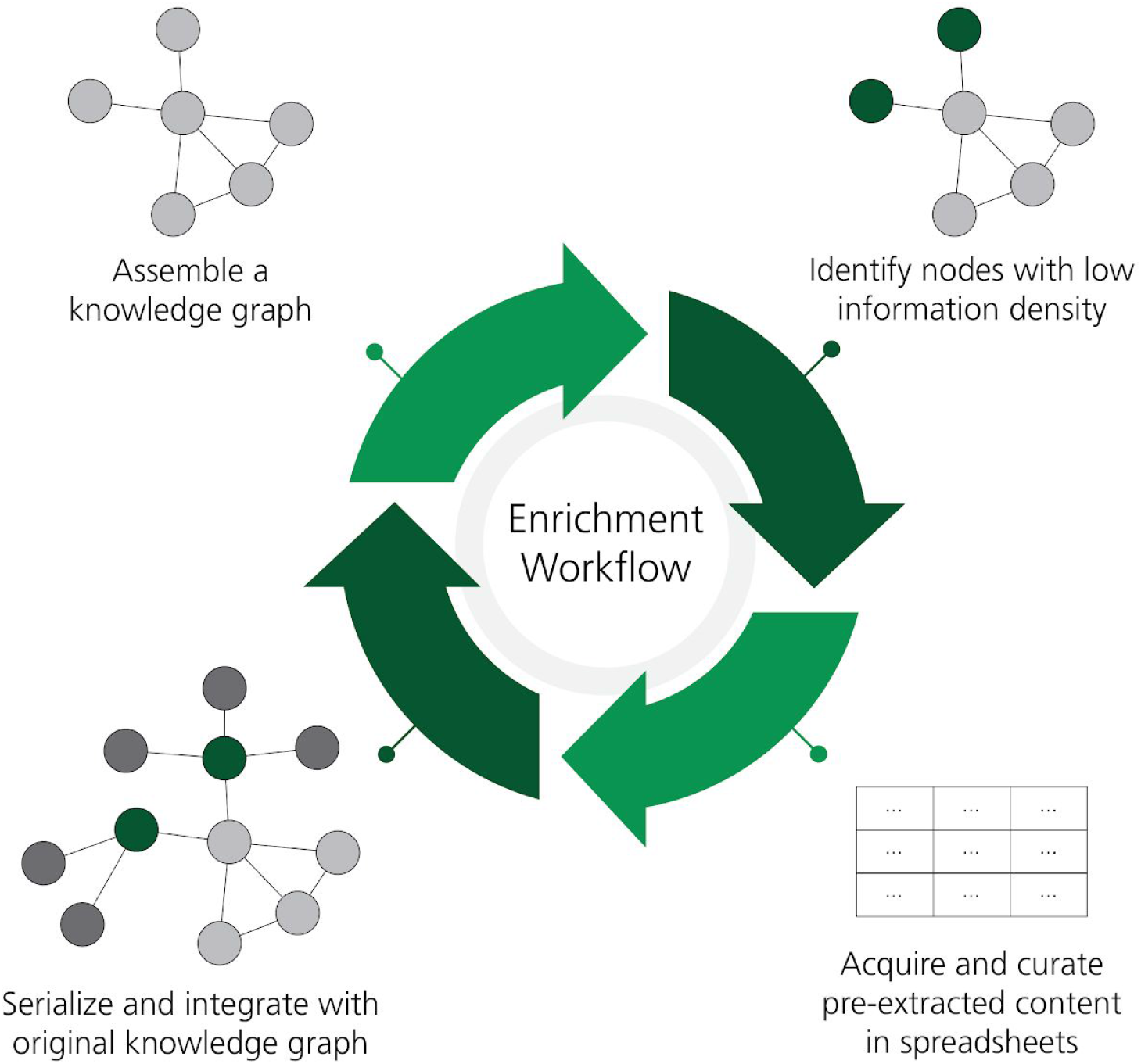
A workflow for the rational enrichment of knowledge graphs. This figure can be found on FigShare at https://doi.org/10.6084/m9.figshare.7642964.v1.

There are two main aspects that are commonly used to formally evaluate a biocuration workflow: the time required to complete the task and quality of the curation compared with a gold standard. To evaluate whether the proposed approach for rational enrichment allows curating a larger amount of statements without compromising the quality, we calculated the average number of minutes required to curate one statement using our proposed workflow and compared it with previous estimates calculated conducting manual curation of BEL statements (Szostak *et al*., 2015; Madan *et al*., 2016) (**Figure 3a**). While the average curation effort was significantly lower than manual curation (2.19 minutes per BEL statement in our workflow vs. 3.2 minutes per BEL statement in manual curation), our calculations included the time used by the curators to annotate the various errors made by the reading system(s). Therefore, if the curation exercise would have exclusively focused on curating BEL statements, the average would have been even lower. Moreover, it is important to note that our proposed approach does not explicitly require the time nor expertise required for corpora generation because the reading systems (e.g., REACH and Sparser) and assembly systems (i.e., INDRA and PyBEL) are applied to all available literature.

**Figure 3.**
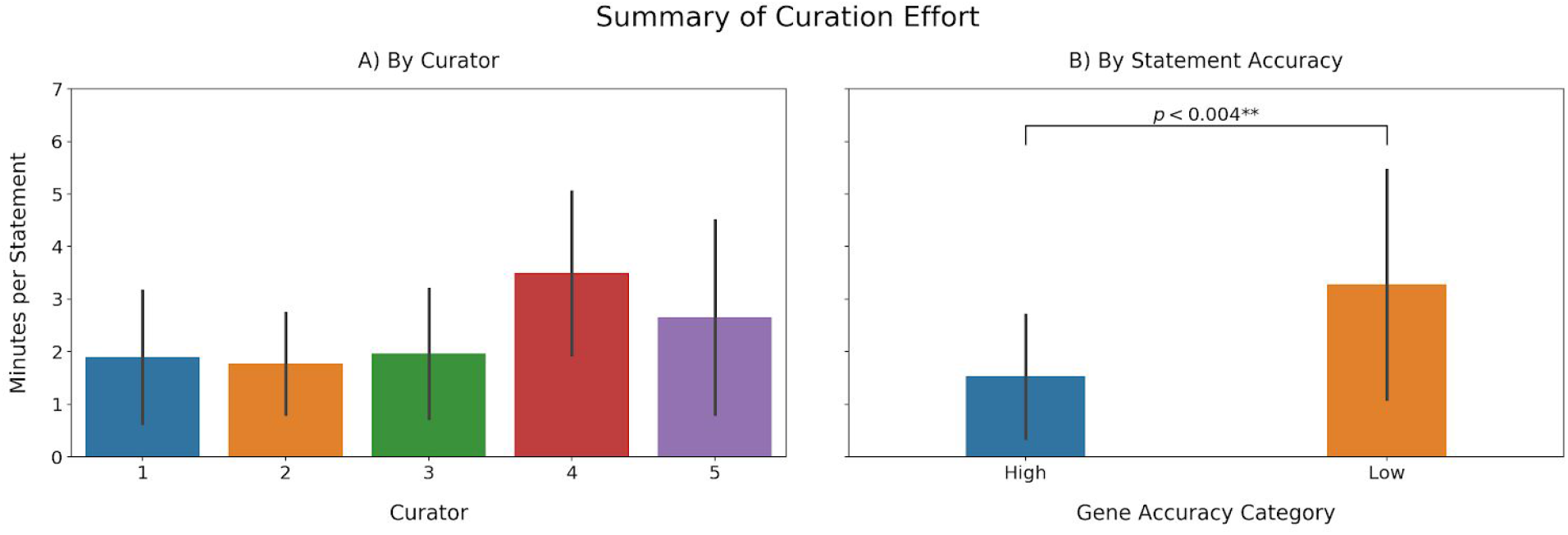
**a**) Recovered BEL statements per minute. Note that the time reported here includes the time invested in annotate the statement as well as INDRA errors. **b**) A comparison of the curation effort between genes for which INDRA had high accuracies (top 20) and genes presenting low accuracies (bottom 20).

Although the amount of time required to curate a certain amount of statements with the proposed approach is lower compared to standard manual curation, the curation effort is also highly variable depending on which gene was curated (**Figure 3a**). To investigate how the curation effort depends on the accuracy of the reader extracting BEL statements, we compared the average curation effort between genes whose statements were accurately and poorly extracted (**Figure 3b**). We observed that the curation effort required to extract statements in genes whose statements were highly accurate (top 20) was significantly less (p < 0.004; Student’s T) than the effort required to curate low accuracy (bottom 20) genes, which effectively took as long as manual curation. We conclude that the high variability associated with the average curation times per curator can be explained by the extra invested time in the genes presenting low recall.

The second aspect we evaluated was the performance in terms of quality. To investigate the direct quality of the BEL statements coming from INDRA, we analyzed the distributions of correct statements before curation observed in each gene (accuracy investigation) (**Figure 4a**). Most of the genes presented accuracies close to the mean accuracy (35.75%) with only a few outliers whose limited number of extracted statements lead to their respective high or low accuracies (see **Supplementary Figure 1**). Furthermore, in accordance with previous research assessing the quality of automatic and manual relation extraction (Rinaldi *et al*., 2016), the accuracies we observed again indicated that BEL statements must be manually curated in order to generate high quality networks. After curation, the distribution of statements that were correct plus statements that were fixed during curation (i.e., excluding statements that were incorrect and could not be fixed) shifted completely to long-tailed distribution with an average of 74.63% BEL statements successfully extracted (**Figure 4b**). The remaining statements (approximately 25%) could either not be coded in BEL nor contained any relevant information about the particular gene.

**Figure 4.**
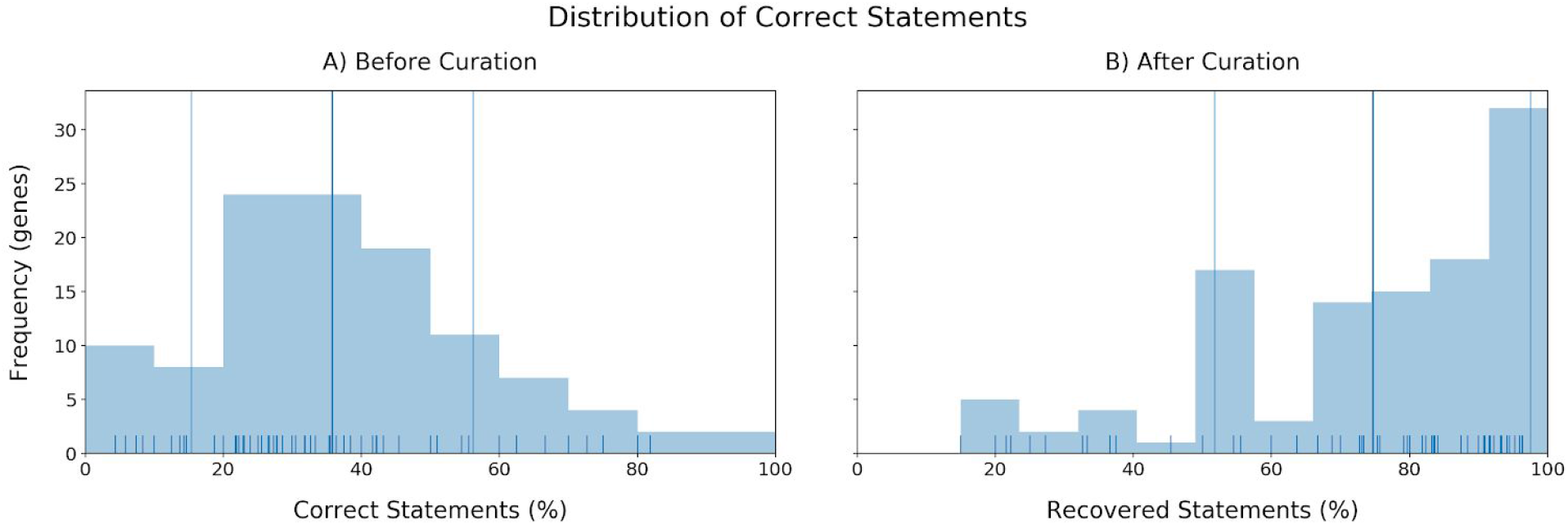
**a**) The distribution of the accuracies in triple identification by INDRA for each gene. X-axis: Correct statements (%). Y-axis: Number of genes (frequency). **b**) Distribution of recovered statements after curation (mean: 74.63%).

While curating the BEL statements, we also annotated the errors made throughout the process of reading, assembly by INDRA, and conversion to BEL by PyBEL in order to identify common mistakes and to assist in the improvement of these three systems. The results showed that the most common error is caused by the name-entity recognition system that identifies the entities participating in the relation (**Figure 5**). Other common errors arose from the improper assignment of the subject and object entities, from evidences that did not actually include relations between the subject and object entities, and statements that were semantically incorrect due to a negation word (e.g., not, no, none, neither, etc.).

**Figure 5.**
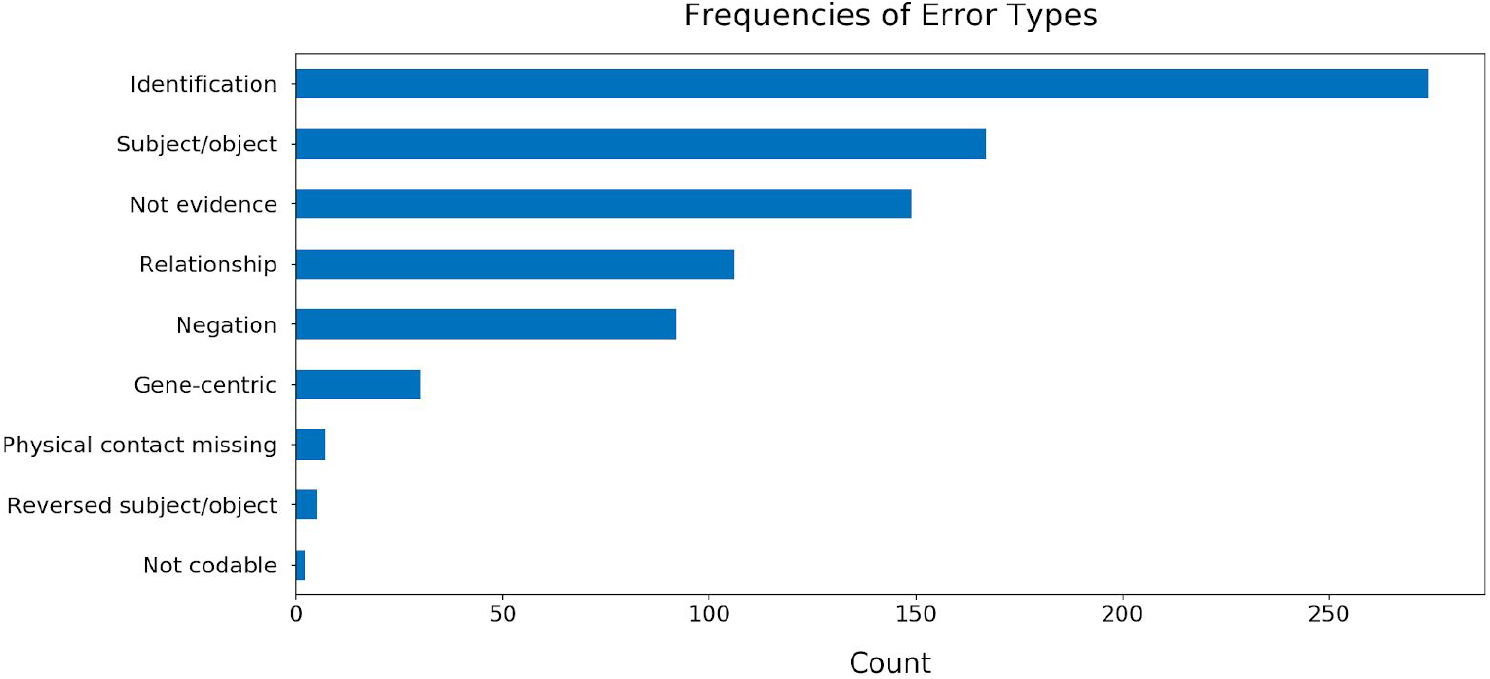
The frequencies of common errors found while curating BEL statements generated from 113 genes. Further details about each error type and the annotation process are available in the guidelines available at https://github.com/pharmacome/curation/blob/master/indra-errors.rst.

The five curators were tasked with tagging interesting examples of the common mistakes that could be used to inform the development of the reading systems (REACH, Sparser, etc.) and the assembly systems (INDRA and PyBEL). Because the authors of this manuscript maintain the INDRA and PyBEL packages, identifying the causes of errors in assembly was relatively straightforward. For example, BEL statements containing biological processes were consistently output using invalid BEL syntax, including the *activity()* function, which is reserved for proteins and other physical entities. We addressed this by updating the previously mentioned *indra.assemblers.PybelAssembler* class. Another error type that was not addressed until after the evaluation was completed was the determination of the role of direct physical interaction in causal relations. INDRA makes use of linguistic cues from the text mining systems along with information from protein-protein interaction databases to determination if a relation involves a physical interaction between proteins, but this information was not incorporated into the *indra.assemblers.PybelAssembler* class. Instead, by default all relations were output using BEL statements implying physical contact: “directly increases” (i.e. increases via contact) and directly decreases (i.e., decreases via contact). This issue has since been fixed. In general, the direct/indirect distinction is difficult to detect automatically in natural language, though it is very important in the generation of mechanistic and mathematical models arising from biological knowledge.

In **Table 3**, we present a small sampling of the errors and corresponding suggestions for improvement in the reading systems. We present a much more thorough enumeration of the errors found in statements for the 113 curated genes in the supplementary information. Besides generating new content quickly, this curation procedure includes information to allow for the evaluation of the automated relation extraction systems and for the proposition of improvements. For example, new groundings can be proposed for entities that were often mismatched. A prominent example was the misidentification of tau (a human protein) and taurine (an amino acid).

**Table 3:** Examples of errors that resulted in suggestions for improvements for the underlying relation extraction systems.

| Gene | Evidence | Issue | Suggestion |
| --- | --- | --- | --- |
| MRC1 | In conclusion, these results suggest that BCR and ABL kinase abrogates MMR activity to inhibit apoptosis and induce mutator phenotype. (Stoklosa *et al.*, 2008) | MRC1, also known as MMR, was confused with Mismatch repair (MMR) | Machine learning methods generating contextual word embeddings could be used to improve the named entity recognition component such as NeuralCoref (https://github.com/huggingface/neuralcoref) |
| TIMP1 | In our work, the restoration of cholesterol efflux capacities from EPA enriched HMDM treated with both the adenylate cyclase activator forskolin and the phosphodiesterase inhibitor IBMX strongly suggests that EPA decreased the ABCA1 mediated cholesterol efflux from HMDM through a PKA dependent pathway. (Fournier *et al.*, 2016) | TIMP1, also known as EPA, was confused with eicosapentaenoic acid (EPA) | Improve the named entity recognition (disambiguation) process, for example, by updating synonym dictionaries in rule-based systems. |
| TRPV1 | Moreover, recently TRPV1 has been demonstrated to be either inhibited or activated by PIP 2. (Morelli *et al.*, 2014) | Only the inhibition relationship was extracted | Rule-based relation extraction systems could be appended with new rules to handle sentences with multiple objects. This and similar examples could be included in the training data for machine learning-based relation extraction. |
| NUMB | This interaction is mediated by the NPXY motif of LNX1 and leads to ubiquitination of Numb by the RING domain of LNX1, thereby targeting Numb to proteasomal degradation. (Young *et al.*, 2005) | The complex sentence structure of “ubiquitination” and “targeting” event were not resolved properly, and the ubiquitination was omitted. | Rule-based systems like REACH that explicitly handle ubiquitination events could be appended with new rules. |
| USF2 | Taken together, the results shown in Figs. 5A, B and C suggest that USF2 stimulates the transcriptional activity of NFκB by enhancing the degradation of IκBα. (Wand *et al.*, 2009) | Relation should be treated as an indirect, rather than direct, increase | Update the INDRA *PybelAssembler* to make use of information about whether a relation is mediated through physical contact. |

Additionally, new rules could be suggested for rule-based systems to avoid issues with the mis-identification of the order of the subject and object as in the example of *“Bak expression was also induced in cells overexpressing the stress-induced transcription factor GADD153, but Bak expression was inhibited in cells expressing an antisense GADD153 construct”* (Lovat *et al*., 2003) whose use of the passive voice may have caused REACH to interpret the statement as “*Bak increased GADD153*.” Ultimately, we believe we can use these examples to provide useful feedback to the developers of the reading systems and improve future extraction.

After applying the re-curation workflow to our selection of knowledge graphs in the NeuroMMSig inventory, we increased the number of nodes from 1188 to 1704 (~1.5x) and edges from 3529 to 5391 (~1.5x). After applying the enrichment workflow, the number of nodes increased to 5850 (~5x) and edges to 23811 (~7x). A more granular summary can be found in **Table 1**. With a 5x increase in nodes, we would expect to see a 10x increase in edges if the new nodes were completely disconnected from the pre-existing nodes in the knowledge graph, which shows that we have been able to maintain the specificity of the knowledge graphs to a reasonable degree. In total, our curators spent 80 hours on the enrichment step to generate 17,002 new BEL statements with an average rate of 3.54 edges per minute. The resulting enriched knowledge graph can be used in reproductions of previous analyses leveraging the NeuroMMSig inventory to assess their robustness, deliver new insights, and improve future analyses when the results are incorporated into a future release of the NeuroMMSig mechanism enrichment server. Additionally, the statements comprise a large training set for future machine learning approaches for text mining.

## Conclusions

We have proposed and applied a generalizable workflow for enriching and updating existing biological knowledge graphs with a focus on the reduction of curation time both in literature triage and in extraction. While its realization involved spreadsheets rather than a *bona fide* curation interface, we believe that it could be adopted by both BEL-specific curation interfaces (e.g., BELIEF, BioDati Studio^1^) and more general biological relation curation interfaces (e.g., NOCTUA^2^, Factoid^3^, WikiPathways (Slenter *et al*., 2017)). Furthermore, INDRA is flexible enough to generate curation sheets for curators familiar with formats other than BEL, such as BioPAX or SBML.

This workflow is by no means the ultimate solution for finding relevant content. Using pre-extracted statements as a stand-in for relevance allows a given knowledge graph to be expanded, but it requires several rounds to find the limits of a given pathway or graph, during which the scope of the curation could be lost. We plan to investigate other methods for identifying relevant content by combining topic modeling with mind maps to not only identify content at the entity level, but on a higher abstraction that allows for capturing of entire areas of biology. These methods could compensate for the simplications that we made to the curation task, such as removing relations containing chemicals, biological processes, and phenotypes. Additionally, they could enable earlier-stage curation that is more focused on achieving reasonable coverage of the available knowledge rather than high granularity enrichment.

Ultimately, as automated relation extraction technologies improve, they will be used to more significantly supplement manual curation efforts. We expect to see many upcoming workflows leveraging these exciting prospects.

## Declarations

### Acknowledgements

We would like to thank Stephan Gebel for his organizational support and Alina Enns and Keerthika Lohanadan for their help in the curation tasks.

### Funding

This work was supported by the Fraunhofer Society under the MAVO project, the Human Brain Pharmacome (https://pharmacome.scai.fraunhofer.de). D.D.F was supported by the EU/EFPIA Innovative Medicines Initiative Joint Undertaking under AETIONOMY [grant number 115568], resources of which are composed of financial contribution from the European Union’s Seventh Framework Programme (FP7/2007-2013) and EFPIA companies in kind contribution.

### Authors’ Contributions

C.T.H. and D.D.F. conceived and designed the study and authored this manuscript. C.T.H., D.D.F., R.A., L.X., S.S., E.W., and K.K. performed curation. J.B., B.G., and P.G. provided data. M.H.A. supervised the project.

### Availability of Data and Materials

The pybel-git Python package that was used to assess syntactic quality is openly available at https://github.com/pybel/pybel-git. All other code and analysis is openly available at https://github.com/bel-enrichment.

### Competing Interests

The authors declare that they have no competing interests.

## Footnotes

1 https://studio.demo.biodati.com

2 http://noctua.berkeleybop.org

3 https://github.com/PathwayCommons/factoid

