## Supplementary Information for "Re-curation and Rational Enrichment of Knowledge Graphs in Biological Expression Language"

| INDRA UUID | B | PMID | Evidence | API | S | P | O |
|---|---|---|---|---|---|---|---|
| c1f943dc-90c5-4612-83a1-89c0d5b32521 | 1.0 | 10782991 | We found that human protein S may indeed activate human Sky but only above physiological plasma concentrations. | reach | p(UP:P27392) | => | act(p(HGNC:TYRO3)) |
| 7b61469c-11c7-497b-bd3a-f7567605d8bd | 0.86 | 12684813 | Finally, EMT is not necessary for the initiation of TGF-beta 1 induced TIF. | reach | p(HGNC:TGFB1) | => | act(p(HGNC:TYRO3)) |
| 7b61469c-11c7-497b-bd3a-f7567605d8bd | 0.86 | 12684813 | Furthermore, macrophages are not important combatants during the early course of TGF-beta 1 induced TIF. | reach | p(HGNC:TGFB1) | => | act(p(HGNC:TYRO3)) |
| c5aeb517-6272-41c5-b17c-cef9bb3ee770 | 0.95 | 18393392 | These observations suggest that Tyro3 RTKs play roles collaboratively in vaginal development, and Mer is more critical, Axl and Tyro3 support the function of Mer. | reach | p(HGNC:TYRO3) | => | act(p(HGNC:MERTK)) |
| 5dc527a0-1c48-48c5-af59-9361b379c026 | 0.86 | 18952193 | This value indicates the TIF is dominant over the TDF, supporting an early, concerted TS. | reach | p(HGNC:TYRO3) | => | act(p(HGNC:TYMS)) |

**Supplementary Table S1.** The first five rows of the pre-curation sheet for hgnc.symbol:TYRO3, produced by the *bel-enrichment* package during the fourth round of curation. The first column, INDRA UUID, represents the INDRA Statement object from which each row was produced and can be used for lookup of other INDRA Evidence objects. The second column, B, represents the belief scores calculated by INDRA based on the confidence in the resources from which its constituent Evidence objects (not shown) come. The third column represents identifier in the PubMed database from which the selected evidence for the statement came. The fourth column, Evidence, corresponds to the selected (normalized) text from the given article that contains the BEL statement written in columns six, seven, and eight. The fifth column, API, represents the reader or knowledge source used to acquire the INDRA Statement. The

full file can be found at
https://github.com/bel-enrichment/results/blob/master/rounds/enrichment-4/TYRO3/TYRO3.bel.tsv.

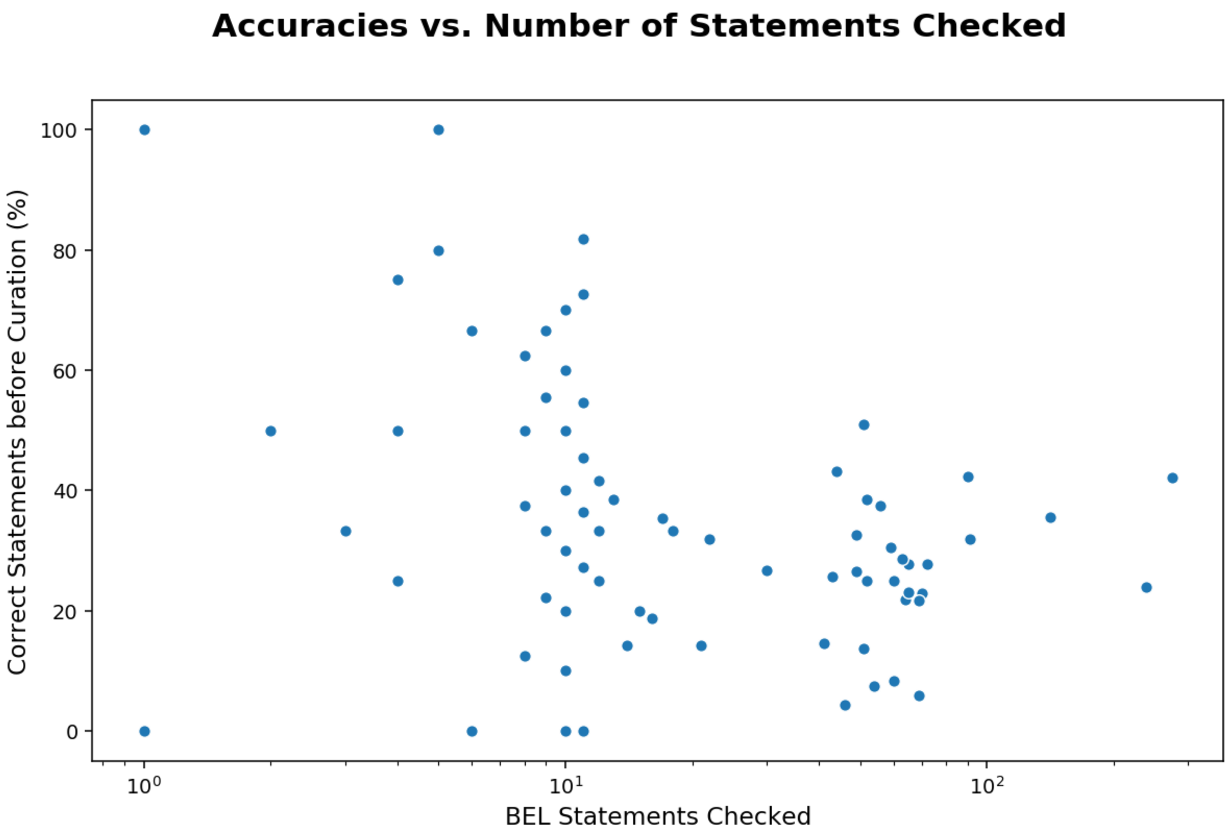


**Supplementary Figure S1.** BEL statements checked vs. correct BEL statements before curation (accuracy of INDRA readers). While most of the genes have an accuracy close to the mean (35.75%), there exist some outliers close to 0 and 100% (dots at the leftmost part of the figure). The figure shows that the reason for these outliers is that they only contained a limited amount of statements.

**Supplementary Text T1.**
The figures and statistics presented in the result section of the manuscript are available in the following GitHub repository: https://github.com/bel-enrichment/results.
